# Novel Role of Copper Transporter CTR1 and Therapeutic Potential of Copper Chelators in Retinal Ischemia-Reperfusion Injury

**DOI:** 10.1101/2025.05.17.654687

**Authors:** Mai Yamamoto, Dipankar Ash, Varadarajan Sudhahar, Syed Adeel H. Zaidi, Modesto A Rojas, Zhimin Xu, Stephanie Kelley, Ruth B. Caldwell, Tohru Fukai, Masuko Ushio-Fukai

**Affiliations:** Vascular Biology Center, Medical College of Georgia at Augusta University, Augusta, GA 30912; Department of Medicine (Cardiology), Medical College of Georgia at Augusta University, Augusta, GA 30912; Department of Pharmacology and Toxicology, Medical College of Georgia at Augusta University, Augusta, GA 30912; Culver Vision Discovery Institute, Medical College of Georgia at Augusta University, Augusta, GA 30912; Department of Cellular Biology & Anatomy, Medical College of Georgia at Augusta University, Augusta, GA 30912; Charlie Norwood Veterans Affairs Medical Center, Augusta GA 30901

## Abstract

**Background:** Retinal ischemia contributes to vision loss in ischemic and diabetic retinopathies through oxidative stress, neurovascular injury, and inflammation. Copper (Cu), while essential, can be toxic in excess and is regulated by Cu transporters such as CTR1. However, the role of CTR1 in ischemic retinopathy remains unclear.

**Methods and Results:** Retinal ischemia-reperfusion (IR) injury was induced by elevating intraocular pressure to 110 mmHg for 40 minutes in the right eye of Ctr1 heterozygous (Ctr1⁺/⁻) and wild-type (WT) mice. In WT mice, IR triggered rapid CTR1 upregulation and increased retinal Cu levels (measured by ICP-MS). IR injury caused retinal ganglion cell loss, inner retinal thinning, vascular degeneration, and apoptosis, all of which were significantly attenuated in Ctr1⁺/⁻ mice. Ctr1⁺/⁻ mice also exhibited reduced microglial (Iba1⁺) and glial cells (GFAP⁺) activation and preserved visual function, as assessed by electroretinography. Mechanistically, IR-induced reactive oxygen species (O₂⁻) production (DHE staining), upregulation of NADPH oxidase components (NOX2, p47phox), and NF-κB activation were markedly suppressed in Ctr1⁺/⁻ mice. Treatment with the Cu chelator tetrathiomolybdate (TTM) similarly reduced retinal thinning, neurovascular damage, apoptosis, gliosis, and oxidative stress after IR injury.

**Conclusions:** CTR1 plays a central role in mediating Cu-dependent oxidative stress, neurovascular degeneration, and inflammation following retinal IR injury. Targeting the CTR1– Cu axis may represent a novel therapeutic strategy for ischemic retinopathy.

## Introduction

Retinal ischemia is a major contributor of visual impairment and plays a key role in several retinal disorders, including glaucoma, diabetic retinopathy, age-related macular degeneration (AMD), and retinopathy of prematurity ^1,2^. These pathologies share common features such as oxidative stress, neurodegeneration, inflammation, glial activation, and vascular damage^1,2^. Experimental models of retinal ischemia/reperfusion (IR) injury, induced by elevating intraocular pressure (IOP), have been widely used to study the underlying mechanisms of neurovascular damage in ischemic retinopathies^3,4,5^. IR injury exacerbates retinal damage by promoting excessive production of reactive oxygen species (ROS), which in turn triggers inflammation, microglial cell activation, apoptosis, resulting in loss of retinal ganglion cell (RGC) neurons, and capillary degeneration, ultimately leading to visual impairment ^6,7,8,9,10^. We previously demonstrated that ROS generated by NADPH oxidase 2 (Nox2) contributes significantly to vascular inflammation in diabetes^11^ and neuronal cell death following retinal IR injury through activation of NFkBp65^12^. However, the precise mechanisms by which IR injury induces ROS production and its link to neurovascular dysfunction remain incompletely understood.

Copper (Cu) is an essential trace element that supports numerous physiological processes, including oxidative phosphorylation, iron metabolism, and the activity of Cu-dependent antioxidant enzymes. However, disruption of Cu homeostasis has been linked to various pathological conditions^13,14,15,16^. Elevated Cu levels have been reported in the serum of patients with diabetic retinopathy^17^ and in the choroid-retinal pigment epithelium (RPE) of donors with AMD^18^. More recently, excess Cu in mitochondria has been shown to trigger *cuproptosis*, a unique form of regulated cell death distinct from apoptosis, ferroptosis, and necroptosis^19^. Given that Cu is both essential and potentially toxic in excess, its intracellular levels are tightly regulated by a network of transporters and chaperones, including Cu uptake transporter 1 (CTR1), Cu chaperones like ATOX1, and Cu-exporting ATPases (ATP7A and ATP7B) ^16,20^.

We recently reported that Cu importer CTR1 expression are markedly increased in the hippocampus of a mouse model of Alzheimer’s disease, as well as in brain endothelial cells (ECs) exposed to amyloid β42, which are associated with elevated Cu levels^21^. Furthermore, our studies demonstrated that CTR1 is critical for vascular endothelial growth factor (VEGF) signaling and angiogenesis in cultured ECs, postnatal retinal vascular development, and mouse models of hindlimb ischemia^22^. Hypoxic conditions also induce CTR1 expression in vascular cells through a HIF1α-dependent mechanism^23^. Of note, elevated CTR1 expression has been detected in the eyes of patients with Eales disease, which is characterized as an occlusive retinal vasculitis^24^, further implicating Cu transport in retinal pathology. Additionally, treatment with the Cu chelator tetrathiomolybdate (TTM) suppresses retinal neovascularization in a mouse model of oxygen-induced retinopathy^25^. Despite these findings, the role of Cu and CTR1 in oxidative stress, neurovascular injury, and neuroinflammation during retinal ischemia/reperfusion injury remains unexplored.

In this study, we demonstrate that the CTR1-Cu axis plays a critical role in promoting neurovascular injury and inflammation in response to retinal IR injury. Using Ctr1^+/-^ mice and Cu chelator TTM, we show that reduced Cu transport attenenuates ROS production and suppresses ROS-dependent NfkB activation in the retina. These findings identify Cu transport proteins, especially CTR1, as a potential therapeutic target for the treatment of ischemic retinal and neurovascular diseases.

## Materials and Methods

### Animal Study

The use of mice was in accordance with the National Institutes of Health Guide for the Care and Use of Laboratory Animals and relevant ethical regulations. The animal protocol used in this study was approved by the institutional Animal Care committee and institutional Biosafety Committee of Augusta University.

### Mouse retinal IR injury model and tetrathiomolybdate (TTM) administration

Both male and female Ctr1^+/−^ heterozygous mice^22^ (Jackson laboratory, 025649) and wild-type (WT) C57BL/6J mice (7–14 weeks old) were anesthetized by intraperitoneal injection of ketamine (87 mg/kg) and xylazine (13 mg/kg). Mice were then subjected to retinal ischemia followed by reperfusion to achieve IR injury as described previously^12,26,27,28^. A 30-gauge needle connected to a raised saline bag was inserted into the anterior chamber of the right eye to raise the intraocular pressure to 110 mmHg (calculated based on the height of the saline bag). Ischemia was induced for 40 min followed by needle removal to allow reperfusion. The left eye served as a sham control since we did not detect differences between contralateral eyes and those from uninjured mice. Animals with a drop in pressure due to saline leakage out of the eye during the procedure were excluded. Mice were deeply anesthetized and sacrificed by transcardial perfusion or cervical dislocation at various time points based on our previous studies and existing literature. For TTM treatment, mice were randomly assigned and gavaged with water (control) or 0.7 mg/day/30g mice TTM daily for 3 weeks. Ceruloplasmin activity was measured in the serum (plasma) of mice with or without TTM treatment using a colorimetric assay based on substrate oxidation (Sigma, Cat# MAK177).

### Optical coherence tomography (OCT)

Mice were placed on the imaging platform of the Phoenix Micron III retinal imaging microscope, supplemented with an OCT imaging device (Phoenix Research Laboratories, Pleasanton, CA). The eyes of mice were dilated with 1% tropicamide (Akron Pharmaceuticals, IL) and genteal gel was applied during imaging to keep the eye moist under ketamine/xylazine anesthesia.

### Retinal thickness measurement

Retinal thickness was studied on retinal fresh frozen sections from mice on 7 days after the IR injury. Cross-sections with optic nerve attachment were prepared (7 *μ*m) followed by H&E staining for morphological observation. Inner nuclear layer thicknesses at three different distances from optic nerve head was determined using ImageJ software and averaged. Averaged retinal thickness was presented as percentage compared with WT sham eyes.

### Electroretinography (ERG) for visual function

A full scotopic electroretinogram (ERG) was recorded on day 7 post-IR as previously described with some modification ^29,30^. Briefly, after 16 h of dark adaptation, mice were anesthetized using a ketamine/xylazine mixture, and the pupils were dilated with 1% tropicamide (Akron Pharmaceuticals, IL) eye drops and genteal gel was applied during imaging to keep the eye moist. Mice were then placed on the electroretinogram system (Diagnosys LLC, Cambridge, UK) with temperature control (37 °C), and dark-adapted ERGs were recorded. Amplitude and implicit times of ERG waveforms were measured at a series of flash intensities (0.001, 0.005, 0.01, 0.1, 0.5, 1 cd.s/m^2^) as previously reported ^29,30^. The a-wave amplitude was measured from the average pretrial base line to the most negative point of the trace, and the b-wave amplitude was measured from that point to the highest positive point.

### Copper (Cu) Measurements

Cu contents were analyzed by inductively coupled plasma mass spectrometry (ICP-MS) using a PlasmaQuad3, as reported previously^21^.

### Dihydroethidium (DHE) staining for detection of superoxide (O_2_^-^)

DHE staining was used to evaluate O_2_^-^ production. Briefly, fresh frozen sections were preincubated with or without SOD-polyethylene glycol (400 U/ml, Sigma-Aldrich, St. Louis, MO, USA) for 30 min, followed by reaction with DHE (2 μM) for 15 min at 37 °C. Sections were washed and mounted. The oxidized DHE, which turns into ethidium bromide upon reacting with superoxide, binds to DNA and fluoresces red. This fluorescence was visualized and captured immediately after staining using a fluorescence microscope (Keyence, BZ-X700) equipped with the appropriate filters (excitation at 488 nm and emission at 610 nm). Fluorescence intensity was analyzed using ImageJ software.

### Immunofluorescence

Cryostat sections with a thickness of 7-12 μm were used for immunofluorescence staining. Fresh frozen sections were employed for NeuN and GFAP staining, while sections fixed in 4% paraformaldehyde (PFA) and embedded in optimal cutting temperature compound (OCT) were used for Ctr1 and Iba1 staining. To prepare the 4% PFA-fixed OCT sections, eyes were fixed overnight in 4% PFA at 4°C. The following day, the eyeballs were thoroughly washed in phosphate-buffered saline (PBS) and then incubated in 15% and 30% sucrose at 4°C for 1 hour for cryoprotection. After sucrose infiltration, the tissues were snap frozen in OCT. Sections were blocked in 10% normal goat serum for 1 hour to prevent non-specific binding. The sections were subsequently incubated with primary antibodies overnight at 4°C (table 1). After primary antibody incubation, sections were washed and incubated with fluorescein-conjugated secondary antibodies for 30 min at room temperature. Following additional washing steps in PBS, slides were covered with a mounting medium containing DAPI (Vectashield; Vector Laboratories, Burlingame, CA, USA) for nuclear staining. Images were acquired using a fluorescence microscope (Keyence, BZ-X700).

**Table 1.** List of antibodies used in IF and WB.

| Antibody | Company | Catalog No, | Dilution |
| --- | --- | --- | --- |
| Anti-CTR1 | Homemade |  | IF (1:100)<br>WB (1:500) |
| Anti-NeuN | Sigma-Aldrich | ABN78 | IF (1:300) |
| Anti-Iba1 | Wako | 019-19741 | IF (1:250) |
| Anti-GFAP | BD Pharmingen | 556330 | IF (1:50) |
| Goat anti-Rabbit IgG (H+L), Alexa Fluor 488 | Thermo Fisher Scientific | A-11008 | IF (1:500) |
| Goat anti-Mouse IgG (H+L), Alexa Fluor 488 | Thermo Fisher Scientific | A-11001 | IF (1:500) |
| Anti-beta Actin | Santa Cruz | sc-47778 | WB (1:1000) |
| Anti-NF-κB p65 | Cell Signaling | 8242 | WB (1:1000) |
| Anti-Phospho-NF-κB p65 (Ser536) | Cell Signaling | 3033 | WB (1:1000) |
| Goat Anti-Mouse IgG (H+L)-HRP Conjugate | Bio-Rad | 1721011 | WB (1:2000) |
| Goat Anti-Rabbit IgG (H + L)-HRP Conjugate | Bio-Rad | 1706515 | WB (1:2000) |

### Tdt-dUTP Terminal Nick-End Labeling (TUNEL) staining for apoptosis

The DNA fragmentation of apoptotic cells was analyzed by terminal dUTP TUNEL assay (In Situ Cell Death Detection Kit, Fluorescein, MilliporeSigma, MA, USA) following the manufacturer’s instructions. The assay measures the fragmented DNA of apoptotic cells by catalytically incorporating fluorescein-12-dUTP at the 3′-OH DNA ends using the enzyme terminal deoxynucleotidyl transferase (TdT). Images were using a fluorescence microscope (Keyence, BZ-X700).

### Retinal vasculature trypsin digestion and counting of acellular capillaries

Eyeballs were isolated on day 14 after IR injury and fixed overnight in 4% PFA. The retinal vasculature was isolated by trypsin digestion as reported ^26,27^. The vasculature was air-dried on silane-coated slides and stained with eosin and hemotoxylin to visualize the acellular capillaries. Acellular capillaries were counted in random fields of the mid-retina using a microscope (Keyence, BZ-X700). The number of acellular capillaries was divided by the field area to get the number of acellular capillaries per 1 mm² of the retina area using ImageJ^26,27^.

### Western Blot Analysis

Retina protein extracts were prepared using RIPA buffer with added protease and phosphatase inhibitors. Protein (25-50 ug) were subjected to SDS-PAGE, transferred to nitrocellulose membranes, and blocked in 5% milk or 2% BSA. Membranes were incubated overnight at 4°C with primary antibodies (table 1), followed by horseradish peroxidase-conjugated secondary antibodies. Bands were visualized using enhanced chemiluminescence and quantified with ImageJ.

### Quantitative Real-Time PCR

Total RNA was extracted from frozen retinal tissue using TRI Reagent (Molecular Research Center Inc. Cincinnati, OH, USA). Reverse transcription was carried out using the high capacity cDNA reverse transcription kit (Applied Biosystems, Waltham, MA, USA) using 2 µg of total RNA. Quantitative PCR was performed with the SYBR Green PCR kit (Qiagen, Germantown, MD, USA) and ABI Prism 7000. The primer sequence for specific genes are shown in Table 2. Samples were run in duplicates to reduce variability. Expression of genes was normalized with HPRT and expressed as fold change of relative mRNA expression compared to control.

**Table 2.**
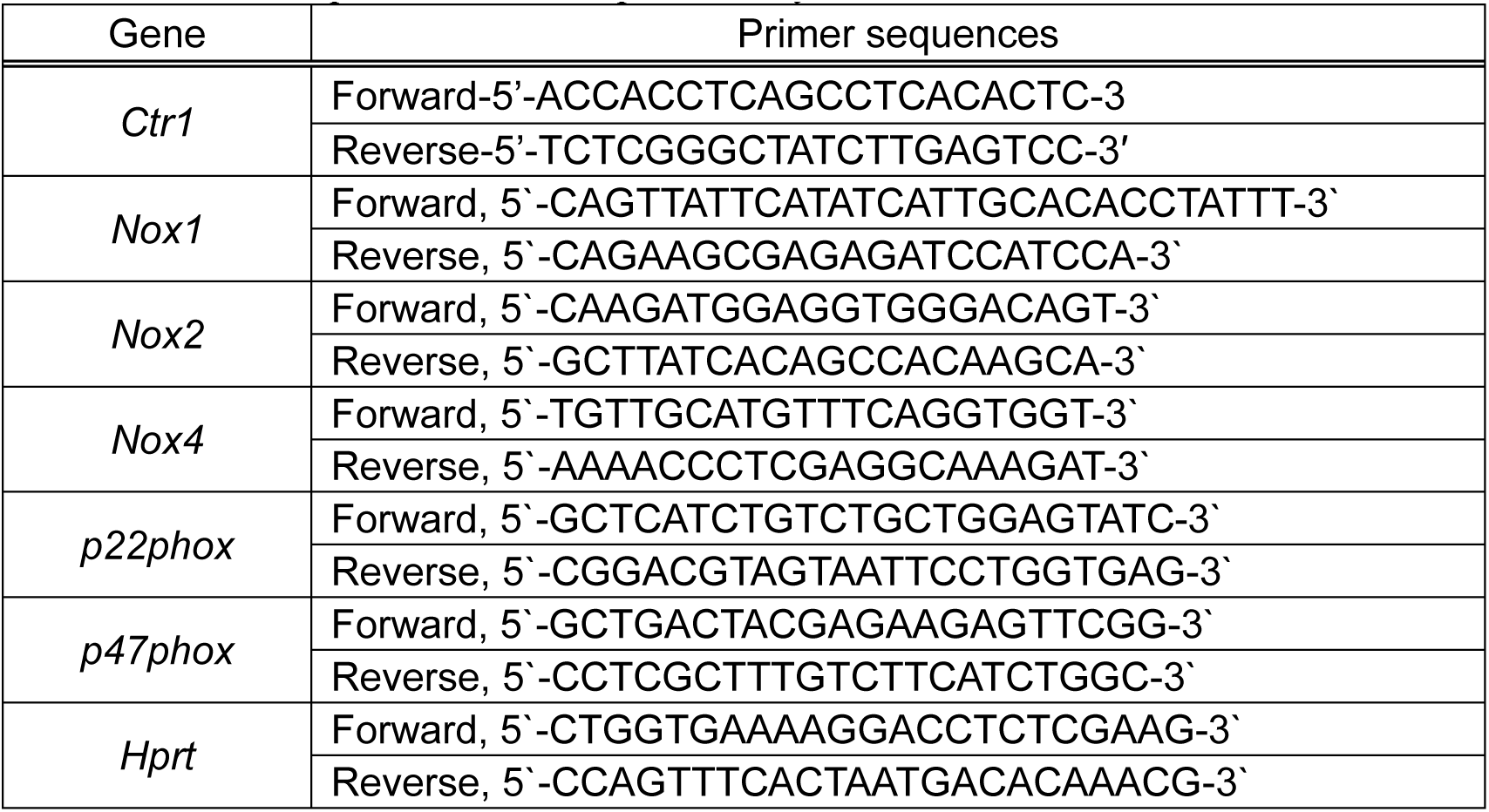
Primer sequences used in qPCR analysis.

### Statistical analysis

Data are presented as mean ± SEM. Statistical tests were performed using Prism v9 (GraphPad Software, San Diego, CA). Data were compared between groups of animals by a 2-tailed unpaired Student *t*-test. One-way ANOVA was applied for multiple comparisons, followed by Sidak’s multiple comparison analysis. The statistical analysis was based on *n*, indicating the number of independent biological samples or mice as described in detail in the respective figure legends. Values of \**p* < 0.05, \*\**p* < 0.01, \*\*\**p* < 0.001 were considered statistically significant.

## Results

### Increased CTR1 expression and Cu Levels in the Retina After IR injury

To investigate the role of the Cu uptake transporter CTR1 in ischemic retinopathy, we utilized a mouse model of retinal IR injury. Mice were subjected to 40 min of ischemia on the right eye followed by reperfusion, while the left eye served as a sham control^12^. Western blot analysis revealed a marked increase in CTR1 protein expression at 3 hours post-injury, which returned to baseline levels by 6 and 24 hours (Figure 1A). Consistent with these findings, immunofluorescence staining showed enhanced CTR1 expression in the outer and inner plexiform layers and the ganglion cell layer at 3 hours post-IR, compared to sham-treated controls (Figure 1B). Importantly, inductively coupled plasma mass spectrometry (ICP-MS) demonstrated a significant increase in retinal Cu content at 3 hours post-injury, with levels normalizing by 6 hours (Figure 1C). In contrast, levels of other trace metals including zinc (Zn), iron (Fe), calcium (Ca), manganese (Mn), and selenium (Se) remained unchanged (Figure 1C and Supplementary Figure 1). These findings suggest that CTR1-mediated Cu uptake is selectively activated during the early phase of retinal IR injury.

**Figure 1.**
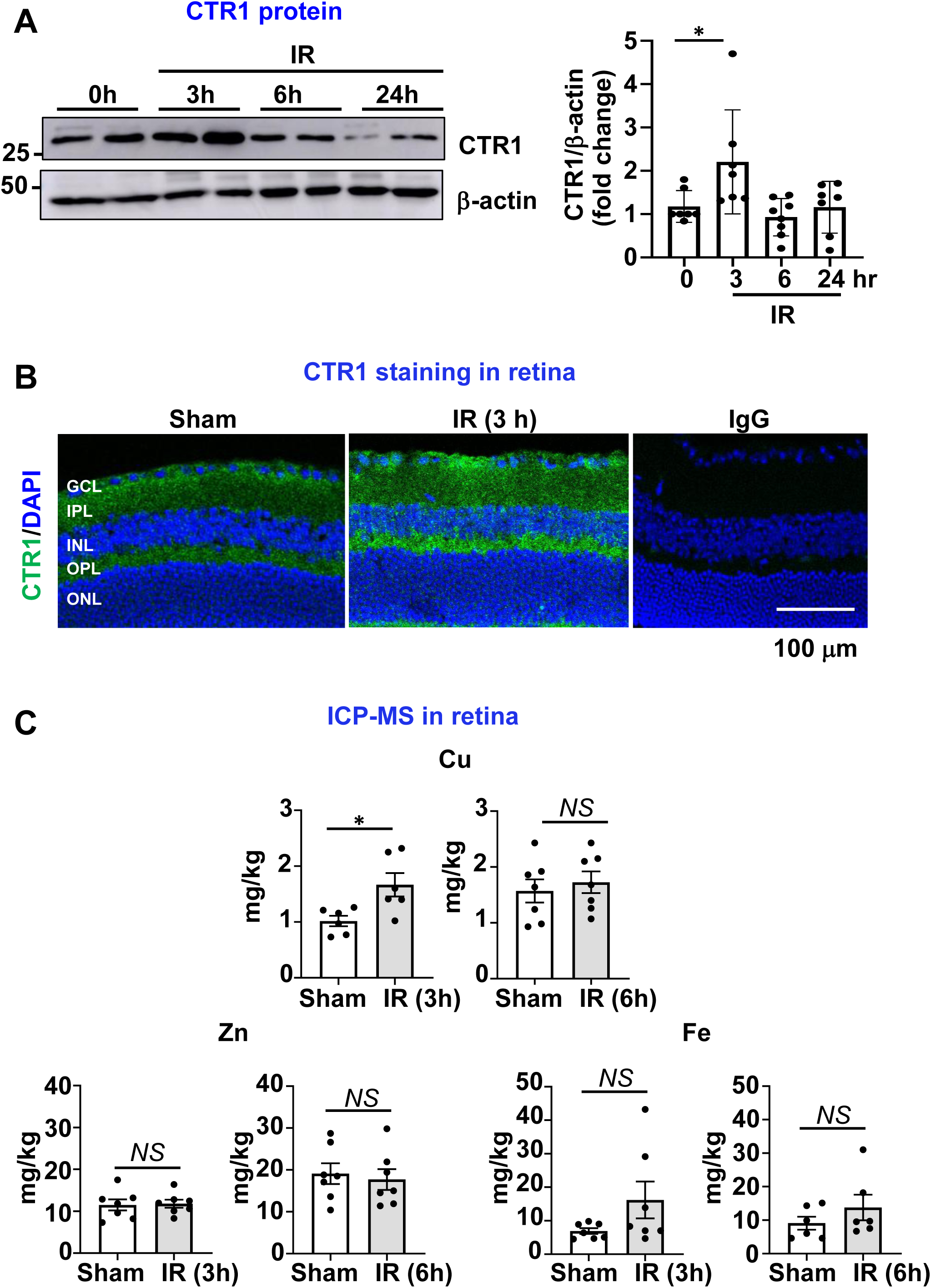
CTR1 protein expression and Cu levels are increased following retinal IR injury in mice. **A**. Western blotting analysis of CTR1 and β-actin (loading control) protein expression in the retinal tissues of WT mice subjected to sham surgery (0 hr) or at 3,6,24 hours after IR injury. Bar graphs show fold changes relative to the sham surgery group, normalized to β-actin. (n=5-8 per group). **B**. Representative immunofluorescence images of retinal tissues stained with CTR1 Ab or IgG (negative control) along with DAPI nuclear staining at 3 hours post-IR injury in WT mice. Scale bar 100 μm. **C.** Copper, zinc and iron contents in the retinal tissues were measured using inductively coupled plasma mass spectrometry (ICP-MS) at sham surgery, 3 and 6 hours post-IR injury. (n=6-7 per group). *p<0.05. NS= not significant.

### Ctr1 hemizygous deletion protects against neurovascular degeneration following IR injury

To investigate the role of endogenous CTR1 in retinal IR injury, we utilized heterozygous Ctr1 knockout mice (Ctr1^⁺/⁻^), as complete knockout of Ctr1 is embryonically lethal^31,32^. Ctr1^+/-^ mice exhibited approximately a 60% reduction in CTR1 mRNA expression (Supplementary Figure 2A). The retinal IR injury is characterized by both neuronal and microvascular degeneration, manifested by retinal ganglion cell (RGC) loss and acellular capillary formation^26^. To evaluate neurodegeneration, we quantified RGC survival in the ganglion cell layer (GCL) using NeuN immunostaining. Seven days after IR injury, WT retinas showed a ∼50% reduction in NeuN-positive RGCs compared to sham controls, whereas Ctr1^⁺/⁻^ retinas exhibited significantly greater neuronal preservation, with only a ∼30% reduction (Figure 2A). To assess microvascular damage, we prepared vascular digests and counted acellular capillaries 14 days post-injury^19,27^. WT retinas displayed a marked increase in acellular capillaries following IR injury, which was significantly reduced in Ctr1^⁺/⁻^ mice (Figure 2B). These results suggest that partial loss of CTR1 confers protection against both neuronal and microvascular degeneration following retinal IR injury.

**Figure 2.**
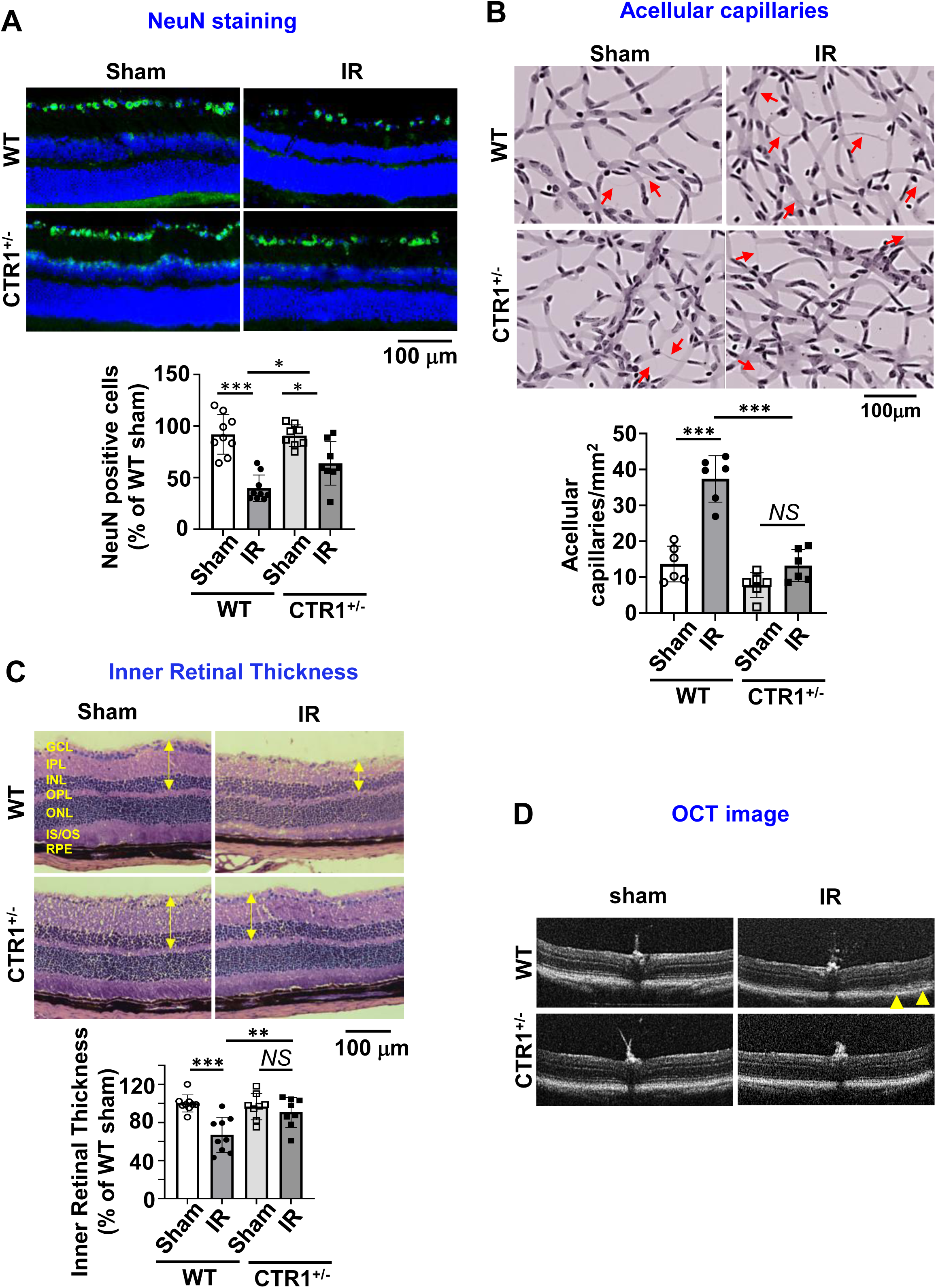
Ctr1 hemizygous deletion protects against neurovascular degeneration following IR injury. **A.** NeuN staining with DAPI nuclear counterstaining in retinal sections from WT and CTR1^+/-^ mice 7 days post-IR injury. Bar graph represents numbers of NeuN-positive cells expressed by the percentage of WT sham. (n=8-9 per group). **B**. Representative images of retinal vascular digests in WT and CTR1^+/-^ mice at 14 days post-IR injury. Red arrows indicate degenerate capillaries. Bar graph represents the number of acellular capillaries per mm^2^ in the retina. (n=6 per group). **C.** Representative H&E-stained retinal sections from WT and CTR1^+/-^ mice 7 days post-IR injury. Bar graph represents the percentage of WT sham. (n=8-9 per group). Abbreviations: GCL, ganglion cell layer; IPL, inner plexiform layer; INL, inner nuclear layer; OPL, outer plexiform layer; ONL, outer nuclear layer; IS/OS, inner segment/outer segment layer; RPE, retinal pigment epithelium layer. **D.** OCT images in the retina of WT retina or CTR1^+/-^ retina at 7 days post-IR injury. Yellow arrow indicates retinal detachment in IR-injured WT retina. *p<0.05, **p<0.01, ***p<0.001, NS= not significant. Scale bar=100 μm.

### Ctr1^+/-^ mice have preserved retinal structure following IR injury

Retinal IR injury primarily affects the inner retinal layers including the GCL, inner plexiform layer (IPL), and inner nuclear layer (INL), resulting in significant thinning of the inner retina ^12,33^. H&E-stained retinal sections revealed a substantial thinning of the (INL in WT IR retinas, with thickness reduced to approximately 60% of sham control levels (Figure 2C). Conversely, Ctr1^+/-^ retinas displayed significantly preserved morphology, with INL thickness at about 90% of sham levels. Optical coherence tomography (OCT) confirmed retinal detachment in WT retinas, while Ctr1^+/-^ retinas remained intact (Figure 2D).

### Ctr1^+/-^ mice have reduced retinal apoptosis and inflammation following IR injury

To explore the mechanisms underlying reduced retinal cell death in Ctr1^⁺/⁻^ mice, we assessed apoptosis and inflammation following IR injury. Retinal neurovascular degeneration is closely linked to apoptosis of neuronal and vascular cells after IR injury^6,34^. TUNEL staining of retinal cryosections revealed a substantial increase in apoptotic cells in WT retinas at 3 days post-injury, whereas Ctr1^⁺/⁻^ retinas showed significantly fewer TUNEL-positive cells, indicating reduced apoptosis (Figure 3A). These findings suggest that reduced CTR1 expression confers protection against IR-induced retinal cell death.

**Figure 3.**
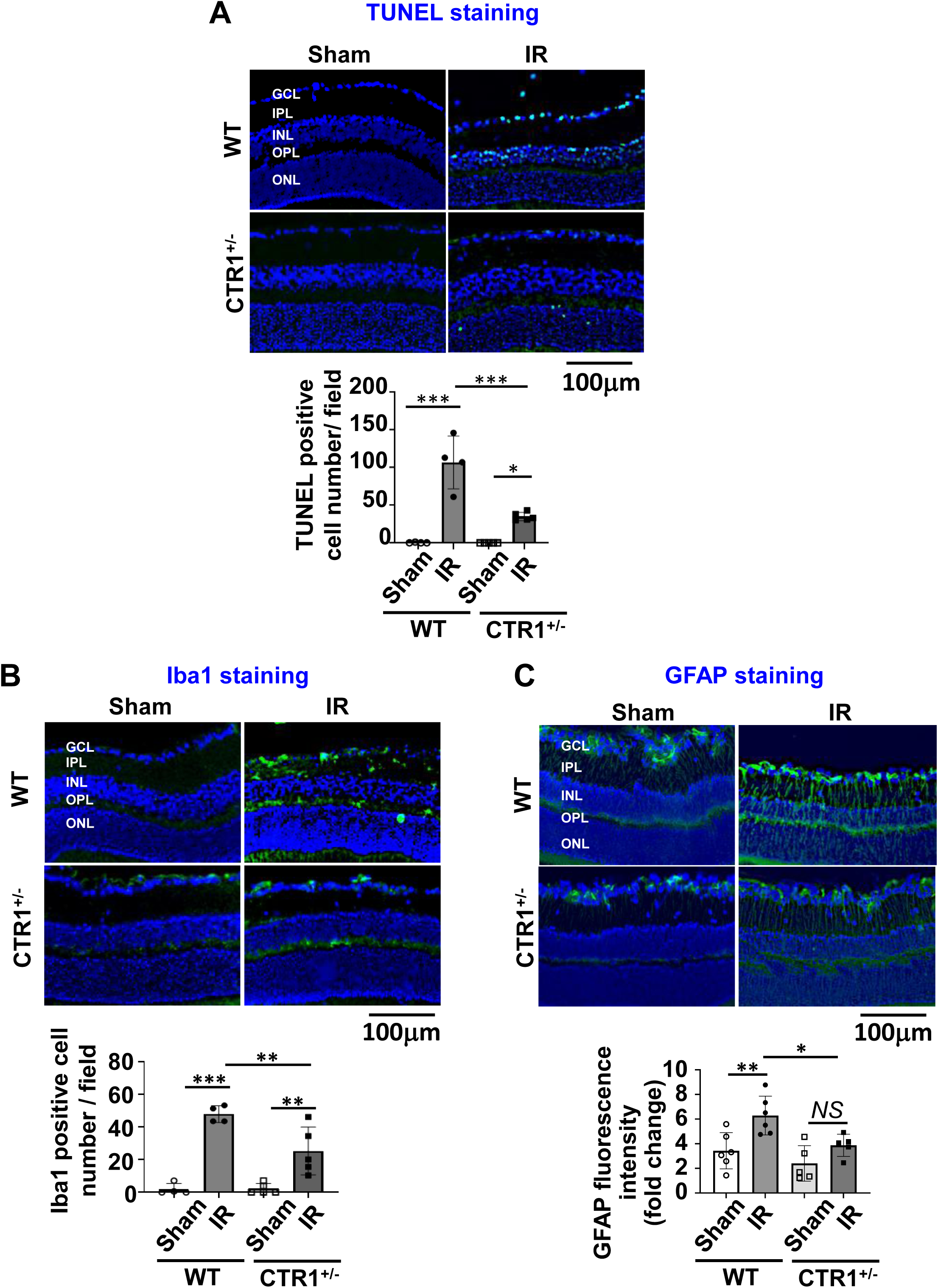
Ctr1^+/-^ mice have reduced retinal apoptosis and inflammation following IR injury. **A.** TUNEL staining with DAPI nuclear counterstaining in retinal sections from WT and CTR1^+/-^ mice 3 days post-IR injury. Bar graph represents numbers of apoptotic cells per retinal section. (n=4-5 per group). **B**. Iba1 staining shows microglia activation, with DAPI nuclear counterstaining, in retinal sections from WT and CTR1^+/-^ mice at 3 days post-IR injury. (n=4-5 per group). **C**. GFAP staining shows Müller cell activation, with DAPI nuclear counterstaining, in retinal sections from WT and CTR1^+/-^ mice at 5 days post-IR injury. (n=5-6 per group). *p<0.05, **p<0.01, ***p<0.001, NS= not significant. Scale bar=100 μm. Abbreviations: GCL, ganglion cell layer; IPL, inner plexiform layer; INL, inner nuclear layer; OPL, outer plexiform layer; ONL, outer nuclear layer.

IR injury is also characterized by robust inflammatory responses^6,34,35^. To evaluate microglial activation, we performed Iba1 immunofluorescence staining. WT retinas showed a significant increase in Iba1-positive microglia at 3 days post-IR injury, whereas this response was markedly suppressed in Ctr1^⁺/⁻^ retinas (Figure 3B). In addition to microglia, activation of glial cells such as astrocytes and Müller cells is a hallmark of IR injury^36,37^. GFAP staining demonstrated strong immunoreactivity in WT IR retinas, with labeling of filamentous astrocytic processes in the nerve fiber layer and radial Müller cell processes extending through the retina. In contrast, Ctr1^⁺/⁻^ IR retinas exhibited significantly reduced GFAP expression in Müller cell processes, suggesting attenuated glial activation (Figure 3C). These results indicate that reduced CTR1 expression mitigates IR-induced inflammation by limiting microglial and Müller cell activation and suppressing apoptosis, thereby providing neurovascular protection in the injured retina.

### Ctr1 hemizygous deletion protects visual function following retinal IR injury

To evaluate the role of CTR1 on retinal function following IR injury, electroretinography (ERG) was performed 7 days post-injury^38^. WT mice exhibited a significant reduction in both a-wave amplitudes, reflecting photoreceptor activity, and b-wave amplitudes, indicating ON bipolar cell function (Figure 4A, B). In contrast, Ctr1^⁺/⁻^ mice showed significantly preserved a- and b-wave responses, indicating reduced functional impairment. These results suggest that partial loss of CTR1 protects against IR-induced visual dysfunction.

**Figure 4.**
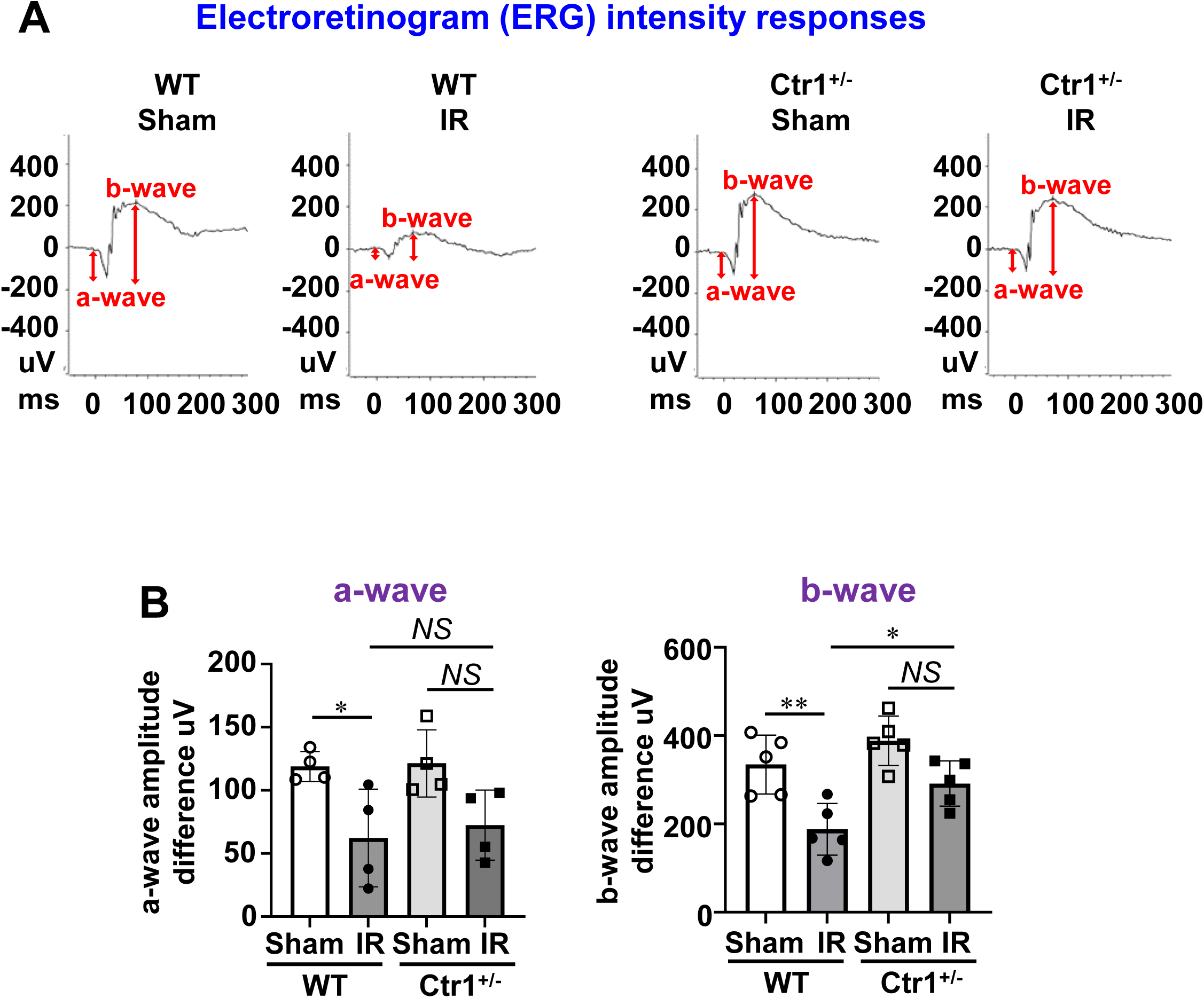
Ctr1^+/-^ mice have preserved visual function following retinal IR injury. **A.** Representative dark-adapted electroretinogram (ERG) intensity showing retinal function by a- and b-waveforms elicited by 1 log cd·s/m^2^ flashes in WT or Ctr1^+/-^ mice at 7 days post-IR injury. **B.** Bar graph represents quantification of a- and b-waveforms amplitudes. (n=4-5 each group). *p<0.05, **p<0.01, NS= not significant.

### Ctr1^+/-^ mice have reduced oxidative stress and suppress NADPH oxidase subunits upregulation following retinal IR injury

Oxidative stress mediated by NADPH oxidase is a critical factor in retinal IR injury. Dihydroethidium (DHE) staining showed a marked increase in superoxide (O₂⁻) production in WT retinas 6 hours after IR injury, which was significantly attenuated in Ctr1^⁺/⁻^ retinas (Figure 5A). We previously demonstrated that IR-induced DHE fluorescence is abolished by PEG-SOD treatment^12^, confirming the specificity for O₂⁻. Quantitative PCR revealed that IR injury upregulated expression of NADPH oxidase subunits *Nox2*, *p22phox*, and *p47phox* in WT retinas, whereas this induction was significantly blunted in Ctr1^⁺/⁻^ mice (Figure 5B). Additionally, western blot analysis showed increased phosphorylation of NF-κB p65, a key downstream effector of ROS signaling, at 3 hours post-injury in WT retinas, which was substantially reduced in Ctr1⁺/⁻ retinas (Figure 5C). These findings indicate that reduced CTR1 expression limits oxidative stress and inflammatory signaling during IR injury by suppressing NADPH oxidase activation and downstream NF-κB signaling.

**Figure 5.**
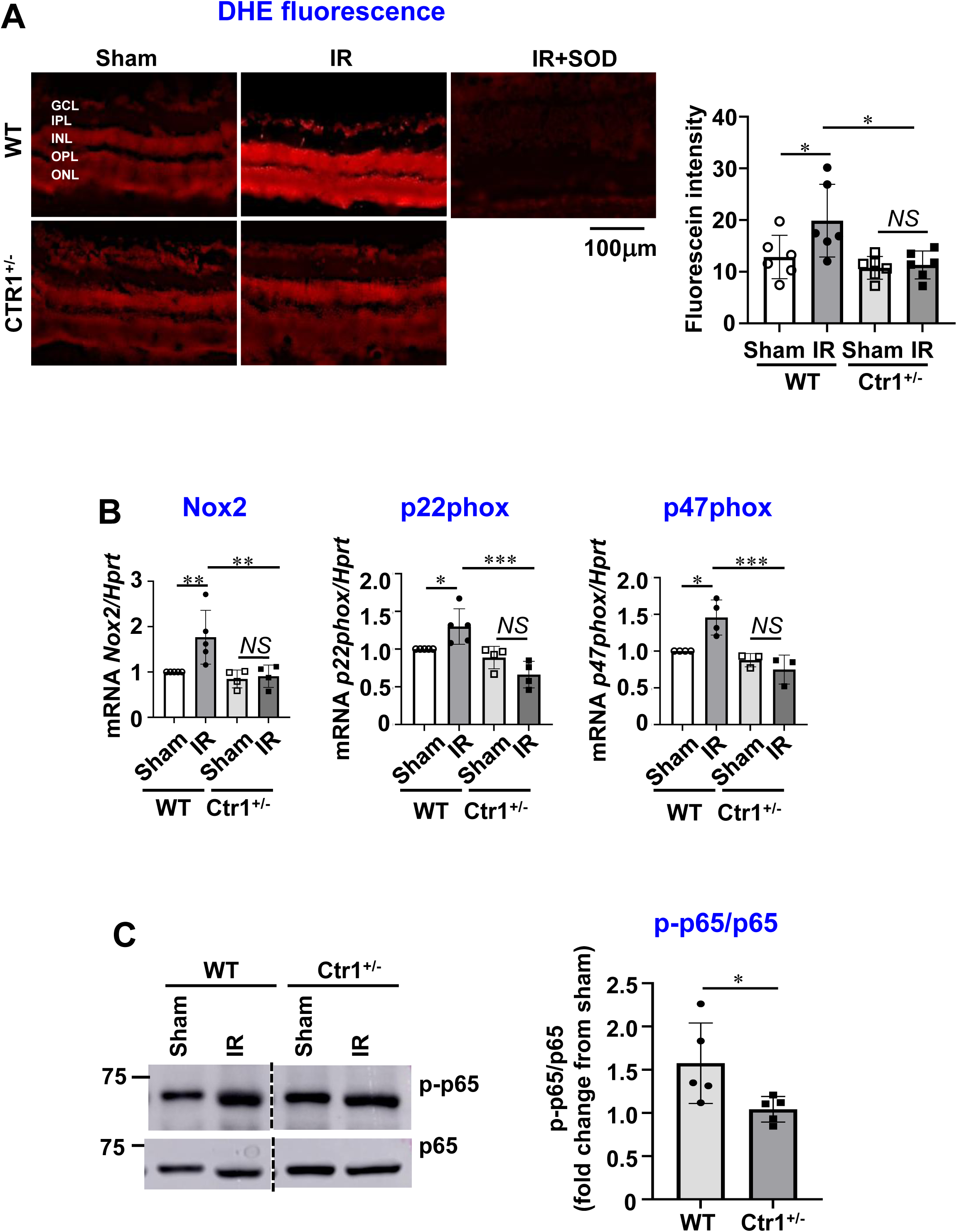
Ctr1^+/-^ mice have reduced oxidative stress and suppress NADPH oxidase subunits upregulation following retinal IR injury. **A**. DHE imaging of O_2_^-^ formation in the retina sections from WT and Ctr1^+/-^ mice at 6 hours after IR injury. Bar graph represents quantification of fluorescence intensity expressed by the average values for each slide (n=6 per group). *p<0.05, ns= not significant. Scale bar=100 μm. Abbreviations: GCL, ganglion cell layer; IPL, inner plexiform layer; INL, inner nuclear layer; OPL, outer plexiform layer; ONL, outer nuclear layer. **B.** qPCR analysis for Nox2, p22phox, and p47phox mRNA expression in retina tissues at 6 hours after IR injury. (n=3-5 per group). **C.** Western blot analysis for phospho-NFkB (p-p65) or total p65 protein expression at 3 hours post-IR injury. Bar graph represents the relative values of each group expressed as a percentage of its respective sham. (n=5 per group). *p<0.05, **p<0.01, ***p<0.001, NS= not significant.

### Cu chelator TTM treatment protects against neurovascular degeneration following retinal IR injury

To determine whether the protective effects observed in Ctr1^+/−^ mice are mediated by Cu, we assessed the impact of TTM, a Cu chelator, in the retinal IR injury model. The efficacy of TTM treatment was confirmed by a reduction in plasma ceruloplasmin activity (Supplementary Figure 2B). TTM-treated mice exhibited preservation of NeuN-positive RGCs in the GCL following IR injury, indicating protection against neuronal loss (Figure 6A). Furthermore, the number of acellular capillaries, a hallmark of microvascular degeneration, was significantly reduced in TTM-treated retinas compared to vehicle controls (Figure 6B). Hematoxylin and eosin staining revealed marked retinal thinning in vehicle-treated animals, which was notably attenuated by TTM treatment (Figure 6C). Consistent with these observations, OCT showed that retinal detachment observed in the vehicle group was not present in TTM-treated retinas (Figure 6D). These results suggest that Cu chelation with TTM confers neurovascular protection during retinal IR injury.

**Figure 6.**
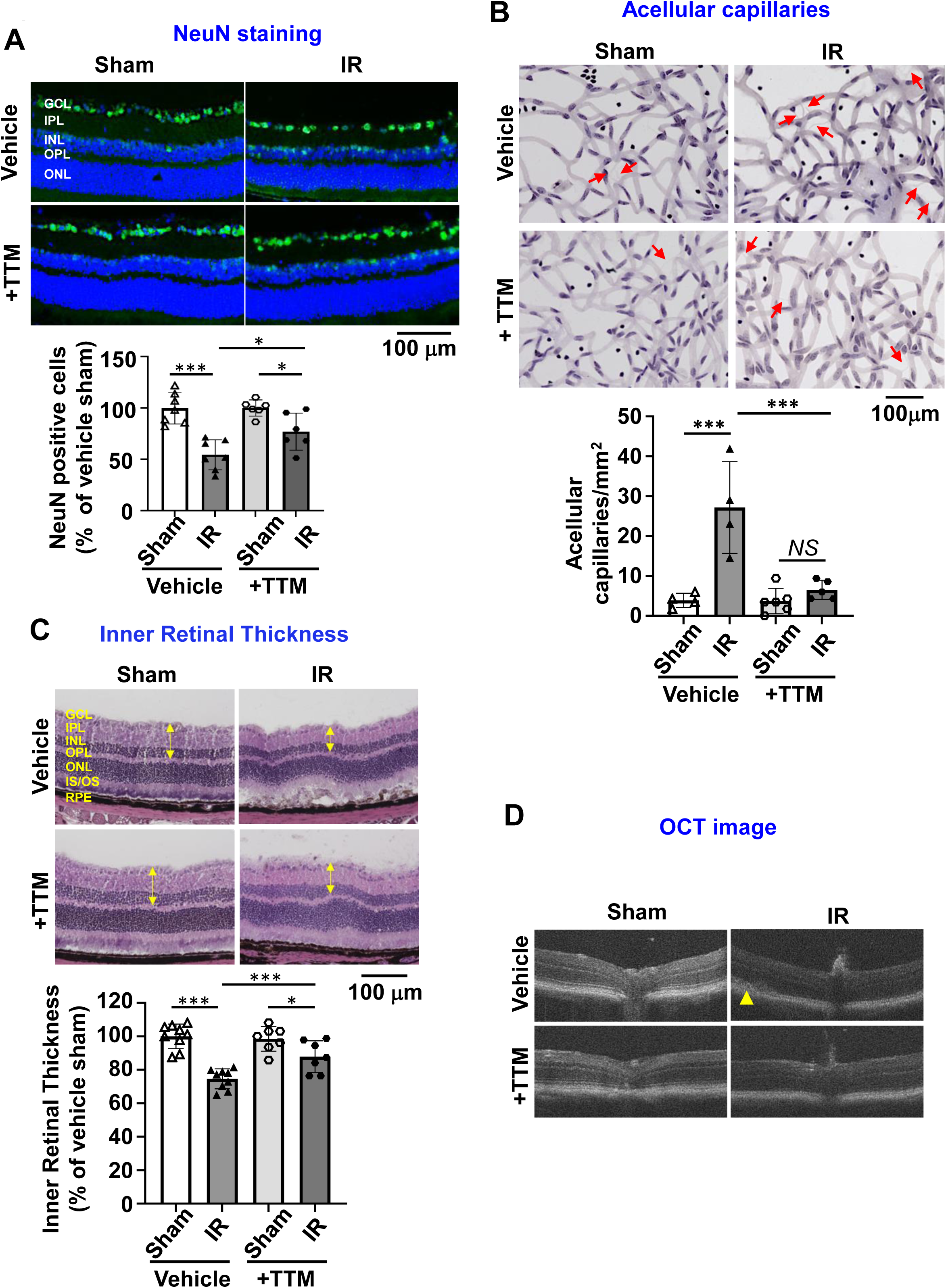
Cu chelator TTM treatment protects against neurovascular degeneration following retinal IR injury. **A.** NeuN staining with DAPI nuclear counterstaining in retinal sections from vehicle-or TTM-treated mice at 7 days post-IR injury. Bar graph represents numbers of NeuN-positive cells expressed by the percentage of vehicle-treated sham. (n=6-7 per group). **B**. Representative images of retinal vascular digests in vehicle- or TTM-treated mice at 14 days post-IR injury. Red arrows indicate degenerate capillaries. Bar graph represents the number of acellular capillaries per mm2 in the retina. (n=4-6 per group). **C.** Representative H&E-stained retinal sections from vehicle-or TTM-treated mice at 7 days post-IR injury. Bar graph represents the percentage of vehicle-treated sham. (n=7-9 per group). Abbreviations: GCL, ganglion cell layer; IPL, inner plexiform layer; INL, inner nuclear layer; OPL, outer plexiform layer; ONL, outer nuclear layer; IS/OS, inner segment/outer segment layer; RPE, retinal pigment epithelium layer. **D.** OCT images in the retina of vehicle- or TTM-treated mice at 7 days post-IR injury. Yellow arrow indicates retinal detachment in IR-injured vehicle-treated mice. *p<0.05, **p<0.01, ***p<0.001, NS= not significant. Scale bar=100 μm.

### Cu chelator TTM treatment reduces retinal apoptosis and inflammation following IR injury

To further investigate the role of Cu in retinal cell death observed in Ctr1^⁺/⁻^ mice after IR injury, we evaluated the impact of TTM treatment. TUNEL staining revealed a significant decrease in apoptotic cell numbers in TTM-treated retinas compared to vehicle-treated controls (Figure 7A).

**Figure 7.**
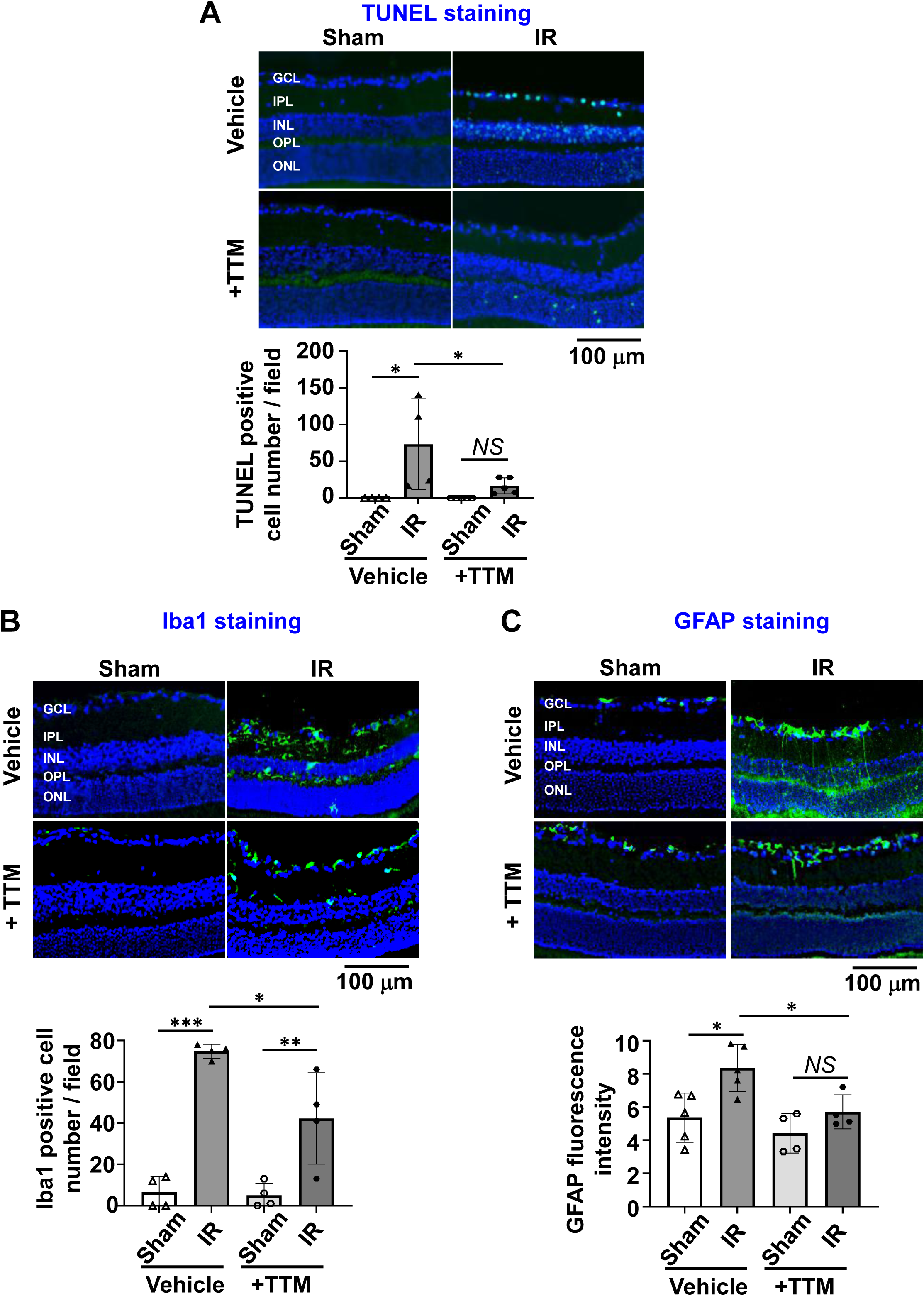
Cu chelator TTM treatment reduces retinal apoptosis and inflammation following IR injury. **A.** TUNEL staining with DAPI nuclear counterstaining in retinal sections from vehicle- or TTM-treated mice at 3 days post-IR injury. Bar graph represents numbers of apoptotic cells per retinal section. (n=4-5 per group). **B**. Iba1staining shows microglia activation, with DAPI nuclear counterstaining, in retinal sections from vehicle- or TTM-treated mice at 3 days post-IR injury. (n=4 per group). **C**. GFAP staining shows Müller cell activation, with DAPI nuclear counterstaining, in retinal sections from vehicle- or TTM-treated mice at 5 days post-IR injury. (n=4-5 per group). (A-C). *p<0.05, **p<0.01, ***p<0.001, NS= not significant. Scale bar=100 μm. Abbreviations: GCL, ganglion cell layer; IPL, inner plexiform layer; INL, inner nuclear layer; OPL, outer plexiform layer; ONL, outer nuclear layer.

To evaluate the role of Cu to IR-induced inflammation, we investigated the effects of TTM on microglial and glial activation following retinal IR injury. Retinas from TTM-treated animals exhibited a significant reduction in IR-induced increase in Iba1 (Figure 7B) and GFAP (Figure 7C) immunoreactivity. These results suggest that Cu chelation by TTM mitigates retinal IR injury by decreasing apoptosis and suppressing activation of microglia and Müller glial cells.

### Cu chelator TTM treatment attenuates oxidative stress following retinal IR Injury

To investigate the mechanisms underlying protective effects of TTM, we evaluated retinal ROS production after IR injury. DHE staining demonstrated that TTM treatment markedly reduced O₂⁻ levels in the retina compared to vehicle-treated controls (Figure 8A). These findings indicate that Cu chelation effectively lowers oxidative stress, supporting the role of Cu in driving ROS generation and contributing to the protective phenotype observed in Ctr1^⁺/⁻^ mice.

**Figure 8.**
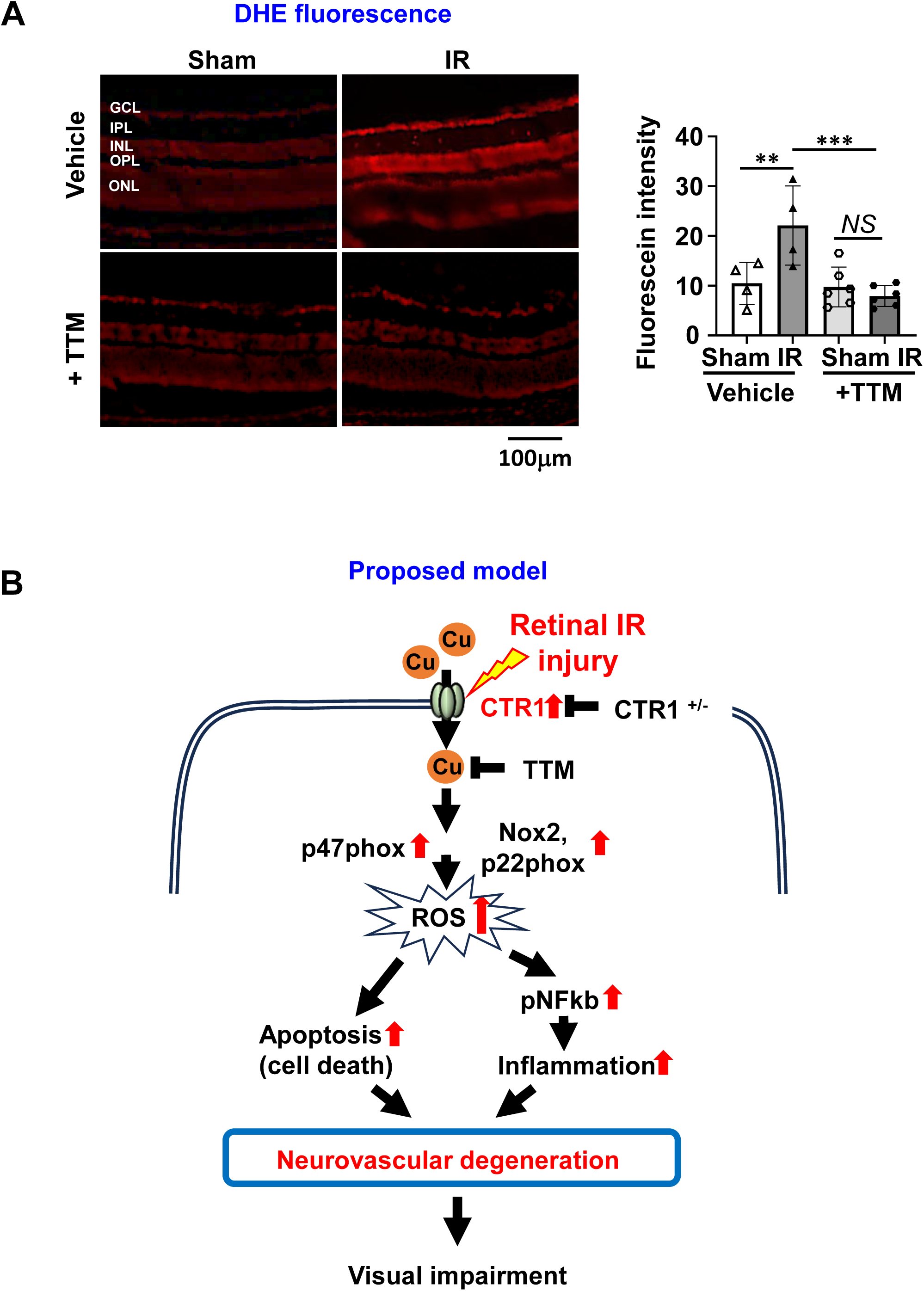
Cu chelator TTM treatment attenuates oxidative stress following retinal IR Injury. **A**. DHE imaging of O_2_^-^ formation in the retina sections from vehicle- or TTM-treated mice at 6 hours after IR injury. Bar graph represents quantification of fluorescence intensity expressed by the average values for each slide. (n=4-6 per group). GCL, ganglion cell layer; IPL, inner plexiform layer; INL, inner nuclear layer; OPL, outer plexiform layer; ONL, outer nuclear layer **p<0.01, ***p<0.001, NS= not significant. Scale bar=100 μm. **B. Proposed model illustrating the role of CTR1-Cu axis in IR-induced inflammation, cell death and neurovascular degeneration leading to visual impairment**. Retinal IR injury triggers a rapid increase in CTR1 expression and Cu levels in the retina, which upregulates the expression of NOX components, leading to elevated ROS production. This promotes the activation of p-NFkB, inflammation, and apoptosis, driving neurovascular degeneration and visual impairment. These IR-induced retinal damages were significantly mitigated in Ctr1^+/-^ mice or through treatment with the Cu chelator, TTM.

## Discussion

Retinal IR injury is marked by oxidative stress, inflammation, and structural damage, including neuronal loss and degeneration of the retinal microvasculature, ultimately leading to visual dysfunction^1,2,39^. Despite these well-documented outcomes, the molecular mechanisms driving this pathology remain incompletely understood. In this study, we identify the copper transporter CTR1 as a key contributor to IR-induced retinal damage. Our findings show that both Ctr1^⁺/⁻^ mice and WT mice treated with the Cu chelator exhibit: (1) reduced neuronal loss and improved survival of retinal ganglion cells, (2) preservation of retinal architecture and prevention of IR-induced thinning, (3) attenuated glial activation and vascular degeneration, and (4) decreased oxidative stress along with preserved visual function. Together, these data support the concept that reducing the CTR1-Cu axis, either genetically or pharmacologically, may represent a viable therapeutic strategy to protect against IR-induced retinal injury and vision loss.

Degeneration of neurons in the GCL is a key feature of retinal injury following IR insult^34,40^. Increased copper levels have been reported in retinal diseases such as diabetic retinopathy and AMD^17,18^, yet the specific contribution of the Cu transporter CTR1 to retinal IR injury has remained unclear. In this study, we identified a transient upregulation of CTR1 and a concurrent rise in retinal Cu levels within 3 hours after IR injury, coinciding with enhanced ROS production. Notably, reducing CTR1 expression in Ctr1^⁺/⁻^ mice or chelating Cu using TTM significantly decreased neuronal apoptosis, as shown by reduced TUNEL staining and preservation of NeuN-positive cells in the GCL. These findings are consistent with prior studies demonstrating the neuroprotective effects of Cu chelation in various ischemic and degenerative models^41^. Furthermore, both Ctr1^⁺/⁻^ and TTM-treated mice maintained retinal structure and showed reduced formation of acellular capillaries, supporting the idea that neuronal loss precedes structural degeneration and microvascular damage^34^. Collectively, our results suggest that targeting the early elevation of CTR1 and Cu accumulation after IR injury may protect against both neuronal and vascular damage, preserving retinal integrity.

Glial activation is an early and critical event in retinal injury and has been closely linked to visual dysfunction in conditions such as IR injury, hypoxia, and diabetes^42,43^. Under physiological conditions, GFAP expression is restricted to astrocytes^42, 44^. In this study, we observed that retinal IR injury induces microglial activation, assessed by Iba1 immunoreactivity as well as GFAP upregulation in Müller cells, a hallmark of glial activation, which were significantly attenuated in both Ctr1^⁺/⁻^ and TTM-treated mice. In addition, IR-induced visual deficits were rescued in Ctr1^⁺/⁻^ mice. These findings underscore the involvement of the CTR1-Cu pathway in driving glial activation and inflammation, which contribute to neurovascular injury and visual impairment following retinal IR injury.

Oxidative stress is a key driver of vascular degeneration, neuronal loss, and inflammation in the ischemic retina. Based on our prior work linking CTR1 to ROS production in brain microvascular ECs^21^, this study demonstrates that both Ctr1^⁺/⁻^ and TTM-treated mice exhibit significantly reduced O_2_^-^ levels, as indicated by diminished DHE staining in the retina. Consistent with previous reports^45^, CTR1 was expressed with particularly strong expression in the GCL and outer plexiform layer, the regions that showed elevated ROS after IR injury. Our findings suggest that the IR-induced increase in retinal Cu may promote the upregulation of NOX2, its membrane-bound regulatory subunit p22phox, and the cytosolic organizer p47phox, collectively enhancing ROS generation. This is consistent with our earlier data showing that NOX2-derived ROS contributes to vascular inflammation in diabetic models^11^ and drives neuronal cell death via NFκB p65 activation following retinal IR injury¹². Mechanistically, we previously showed that the Cu chaperone Atox1 senses elevated Cu levels and then translocates to the nucleus, where it acts as a Cu-dependent transcription factor to induce p47phox expression in response to TNFα stimulation^46^. Moreover, IR-induced ROS activates NFκB, a redox-sensitive transcription factor known to exacerbate inflammation and neuronal damage in the retina^12^. NFκB not only directly increases Nox2 expression^47^ but also indirectly upregulates p22phox, contributing to further NOX2 activation. These findings support the existence of a positive feedback loop in which ROS generated via the CTR1-Cu-Atox1-p47phox axis activates NFκB, which in turn enhances NOX2 activity, promoting oxidative stress, inflammation, and neurovascular degeneration in retinal IR injury.

This study highlights the therapeutic potential of Cu chelation in ischemic retinopathy, demonstrating that TTM treatment preserves retinal architecture, reduces neuronal loss, and attenuates both glial activation and neurovascular damage by suppressing oxidative stress after retinal IR injury. These findings align with previous studies reporting that TTM confers protection in various ischemic models including myocardial infarction, stroke, and liver ischemia by reducing infarct size, tissue injury, and inflammation^48,49,50^. Future research should explore intravitreal delivery strategies for TTM and other Cu chelators to improve therapeutic efficacy in preserving vision and more effectively limiting inflammation and retinal injury.

In summary, our findings underscore the critical role of CTR1 and Cu in regulating oxidative stress, inflammation, and neurovascular degeneration during retinal IR injury, pointing to promising therapeutic opportunities. Cu chelation approaches, such as TTM treatment, may also hold potential for broader application in other ischemic and neurodegenerative disorders linked to oxidative stress and Cu imbalance. Future studies should evaluate the long-term impact of CTR1 modulation and Cu chelation on visual function and examine their possible synergy with current treatment strategies. Additionally, deeper investigation into the molecular pathways through which Cu interacts with other cellular signaling networks may further elucidate its role in retinal pathology. By identifying the CTR1-Cu axis as a viable therapeutic target, this work lays the groundwork for advancing novel treatments for ischemic retinal and neurovascular conditions.

## Acknowledgements

None.

## Author contribution

M.U.-F., T.F., and M.Y., designed the study; M.Y. (major), D.A., V.S., performed/assisted research; M.U.-F., T.F., M.Y., analyzed data; S.A.Z., M.A.R., Z.X., provided inputs and edited manuscript; R.B.C., provided inputs on experimental design and data interpretation, and edited manuscript; M.M. and S.K. performed mouse genotyping; T.F., M.U.-F. and M.Y., wrote the manuscript. All of the authors reviewed the manuscript. We acknowledge the assistance of Dr. Martina Ralle, Director of USR Elemental Analysis Core at OHSU (partial support from NIH (S10OD028492) for measuring metals levels using ICP-MS.

## Data and materials availability

All data necessary to evaluate the conclusions in this paper are present either in the main text or the supplementary materials. Additional data supporting the findings of this study are available from the corresponding author upon reasonable request.

---

**Supplemental Figure 1.**
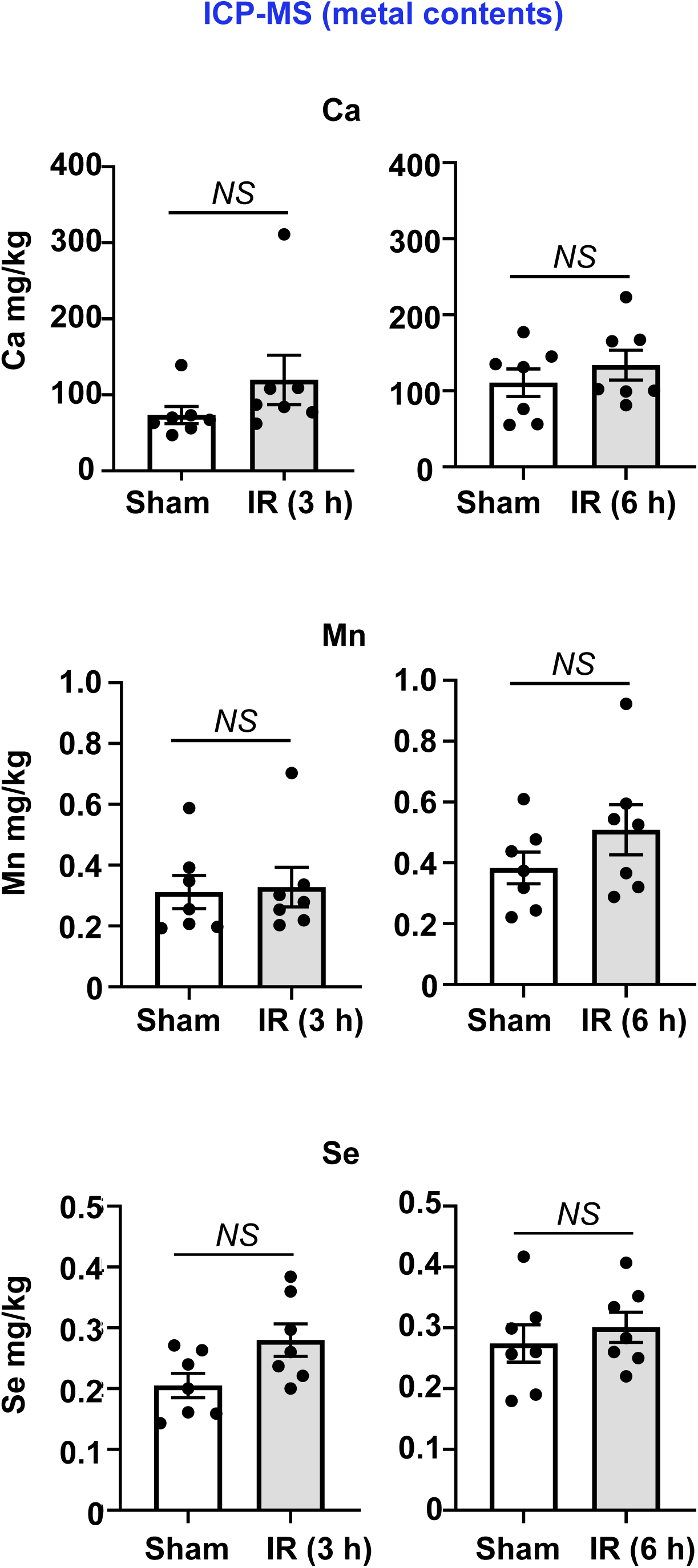
Inductively coupled plasma mass spectrometry (ICP-MS) analysis for Calcium (Ca), manganese (Mn), selenium (Se) contents in the retinal tissues of WT mice subjected to sham surgery or at sham, 3 and 6 hours after IR injury. (n=7 per group). NS: not significant.

**Supplemental Figure 2.**
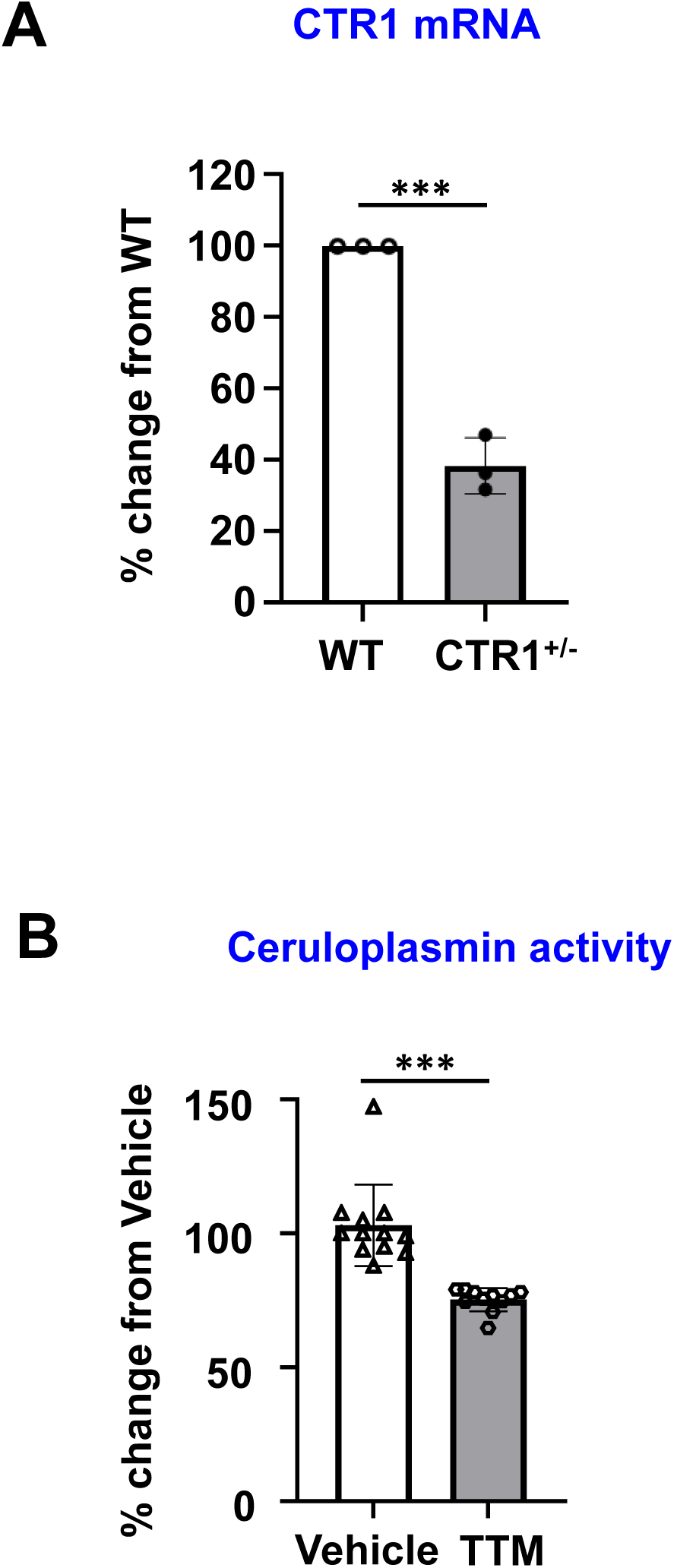
**A.** qPCR analysis for CTR1 mRNA expression in the retina tissues from WT or Ctr1^+/-^ mice. Bar graph represents the % fold change from WT retina. (n=3 per group). **B.** TTM treatment decreased ceruloplasmin activity levels in plasm of WT mice. Bar graph represents the % fold change from WT treated with vehicle. (n=11-12 per group). ***p<0.001.

